# ITGB6 inhibition stimulates anti-tumor responses in immunocompetent mouse models of head & neck squamous cell carcinoma and pancreatic adenocarcinoma

**DOI:** 10.1101/2024.04.18.590156

**Authors:** William J. MacDonald, Praveen R. Srinivasan, Maximilian Pinho-Schwermann, Shengliang Zhang, Vida Tajiknia, Connor Purcell, Jillian Strandberg, Wafik S. El-Deiry

## Abstract

ITGB6, the gene encoding the β6 subunit of integrin αvβ6, is a potent prognostic marker across multiple cancer types. As a major activator of latent TGFβ, αvβ6, and consequently, ITGB6, has considerable therapeutic implications due to the immunosuppressive effect that activated TGFβ has on the tumor microenvironment. The present study identifies ITGB6 as a potent target for inducing an immune-mediated anti-tumor response. ITGB6 is highly upregulated in various squamous cell carcinomas and pancreatic adenocarcinomas, allowing it to disrupt tumor-immune cell signaling, while avoiding the widespread side-effects of systemic TGFβ inhibition. Genetic knockout of ITGB6 in heterotopically injected head and neck squamous cell carcinoma and pancreatic adenocarcinoma cell lines showed markedly reduced tumor progression in immunocompetent mice. Additionally, co-cultures of human squamous cell carcinoma cell lines and human T-cells showed increased T-cell killing upon cancer cell ITGB6 inhibition. Colony formation experiments give further evidence that the reduction in tumor growth observed upon ITGB6 inhibition *in vivo* is through immunological clearance of cancer cells and not merely through intrinsic factors. Analysis of The Cancer Genome Atlas (TCGA) revealed not only the high prognostic value of ITGB6 on overall survival but also that high ITGB6 expression in patients is often associated with an inferior response to α-PD-1 and α-PD-L1 immune checkpoint blockade. The potent anti-tumor immune response observed both *in vitro* and *in vivo* upon ITGB6 inhibition, combined with our analysis of RNA-seq data from immune checkpoint blockade-treated patients, encourages the development of ITGB6 blockade and immunotherapy combination regimes. Further pre-clinical studies will serve to facilitate the translation of our findings into therapeutic clinical trials of combination therapies for treating immunotherapy-resistant cancers.

**Visual Abstract:** 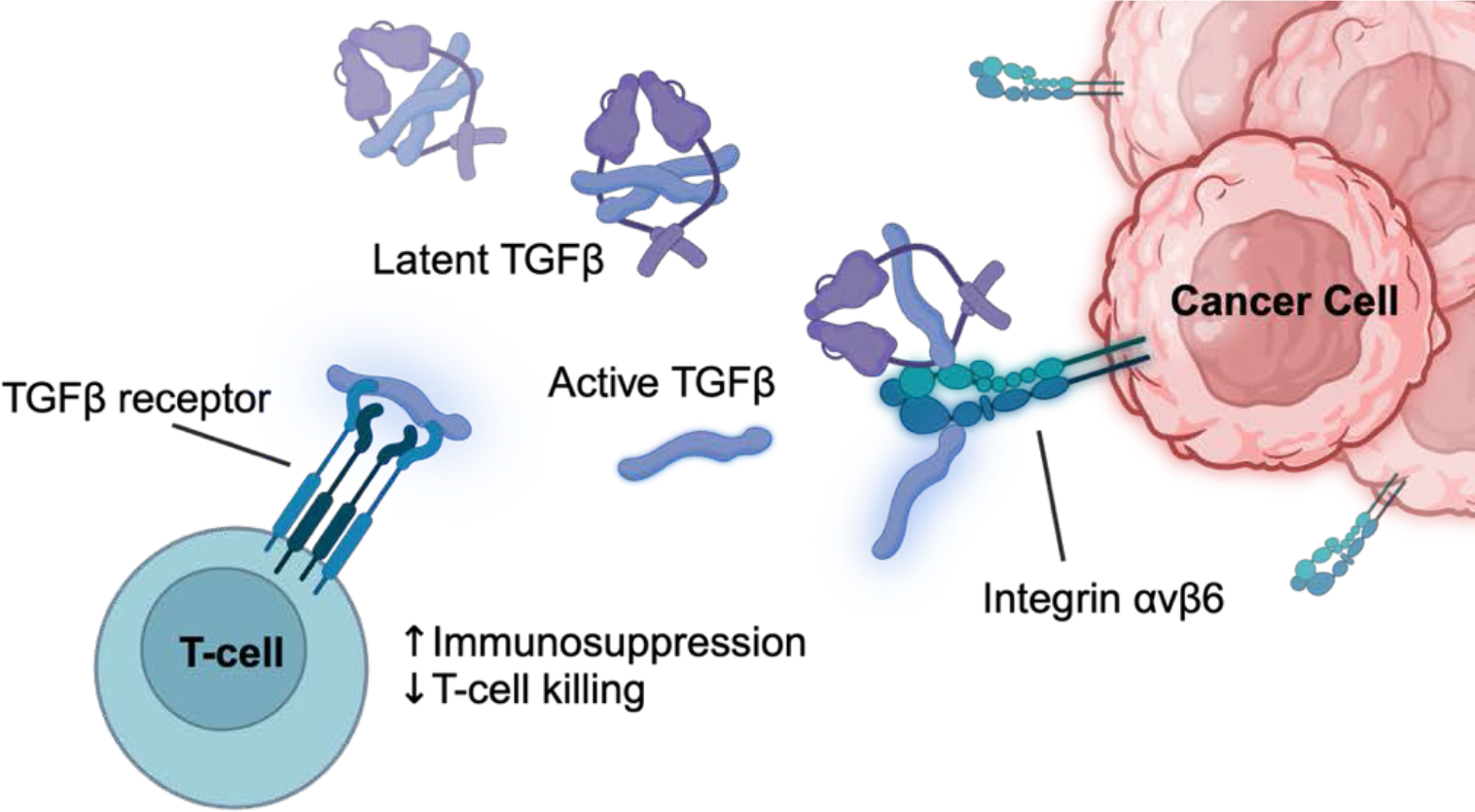

## Introduction

Immunotherapeutic agents that harness the tumor protective power of the immune system have given new hope to treating cancer types often characterized by treatment resistance and poor prognosis. Immune checkpoint blockade therapy against PD-1 and PD-L1, for instance, has proven effective in a variety of cancers. Lung cancer, metastatic melanoma, genitourinary cancers, and head and neck cancer have shown some susceptibility to PD-1/PD-L1 inhibitors, with effective response rates of 29.03%, 26.91%, 20.66%, and 12.15% respectively [1]. Responders will generally experience a duration of response of around one year.

Unfortunately, the majority of patients with these tumor types will see no response to immune checkpoint blockade (ICB) therapy. Among factors such as PD-L1 expression, neo-antigen burden, and immune cell infiltration, one of the main factors that has been observed to influence response to ICB is the cytokines profile of the tumor microenvironment (TME). The cytokine TGFβ, in particular, is associated with a dysfunctional immune response in the TME [2].

TGFβ is a widely implicated cytokine that has disparate effects across cell types. However, in the tumor microenvironment, increased TGFβ signaling has an immunosuppressive effect on T-cells and natural killer cells, resulting in cancer evading the immune system [3]. Decreased local TGFβ around tumors correlates with higher T-cell cytotoxicity [4]. Consequently, TGFβ decreases the effectiveness of immune checkpoint blockade therapy by reducing the infiltration of immune cells into the tumor. Additionally, TGFβ has been shown to drive epithelial-mesenchymal transition (EMT) forcing tumors to a more invasive phenotype with greater metastatic potential [5].

Consequently, The TGFβ pathway is one of the most abundantly mutated signaling cascades in cancer [6]. In normal tissue and early carcinogenesis, TGFβ has a growth-suppressive effect [7]. However, as a malignancy progresses, mutation of the TGFβ pathway, especially the SMAD target genes, can lead to loss of the negative feedback loop governing TGFβ production. These mutations not only cause increased secretion of TGFβ into the TME but also render the cancer cells themselves insensitive to the growth-inhibiting effects of TGFβ. The ability of cancer cells to use TGFβ’s potential to quell a tumor immune response while insulating themselves from the cytokine’s anti-proliferative effects makes for a potent evolutionary strategy that is harnessed by many cancers.

Inhibition of TGFβ has frequently been shown to be effective against cancer via immunological mechanisms. Studies showing sensitization of otherwise ICB-resistant tumors due to TGFβ blockade make clear the immense potential that TGFβ inhibition has for expanding immunotherapy to a far wider cohort of patients [8]. There have been many therapeutic attempts to combat this immune evasion strategy in cancer. However, decades of preclinical success have failed to translate into the clinic as there is currently not a single FDA-approved TGFβ inhibitor for cancer. These trials have often fallen short due to inconsistent efficacy [9]. Additionally, TGFβ is a highly promiscuous cytokine that is widely implicated in normal physiological processes. Therefore, the dose limitation that is necessary to avoid unacceptable off-target effects may make the necessary therapeutic window unattainable.

To avoid the toxicity that comes from inhibiting TGFβ, manipulation of other pathway components is under investigation [10]. TGFβ is predominantly found sequestered in the extracellular matrix (ECM), where it is present in its latent form. Latent TGFβ must be activated before it can be an effector in the TME. TGFβ is activated by various ligands that are differentially expressed across tissue types. The integrin αvβ6, is of particular interest, as it is not only one of the main activators of TGFβ, but it is also expressed at high levels in certain cancer types [11].

αvβ6 is one of 24 transmembrane integrin receptor proteins that facilitate molecular communication between cells and with the ECM. Besides mechanically anchoring cells into the ECM, thus facilitating cell adhesion, integrins serve as mediators of various intracellular and extracellular physiological processes by inducing intracellular transcription factors or activating extracellular molecules for paracrine signaling. As heterodimeric proteins, integrins have an α and a β subunit. Various combinations of the presently identified α and β subunits make up the 24-member integrin family. Integrin αvβ6 plays a powerful and widespread role in cancer biology. αvβ6 is comprised of the αv and β6 subunits, which are encoded by the genes *ITGAV* and *ITGB6 genes*, respectively. While αv complexes with several other β subunits, β6 shows no affinity for other α subunits. Therefore, β6 can be considered the “rate-limiting” subunit for the formation of αvβ6. Consequently, *ITGB6* is generally regarded to be the principal gene responsible for the formation and functioning of αvβ6 [12].

The main extracellular function of αvβ6 is the activation of transforming growth factor-β1 (TGFβ1), which is the member of the TGFβ family most relevant to TME immunosuppression [13]. Integrin αvβ6 contains on its β6 subunit a region that binds the tripeptide arginine-glycine-aspartic acid (RGD) motif found on various cell-surface and extracellular proteins. One of these ligands is the latency-associated protein that forms the latent complex of TGFβ1 that is commonly sequestered in the ECM [14]. Upon binding to αvβ6, active TGFβ1 is released and can function as a cytokine in an autocrine or paracrine manner.

Since αvβ6 is preferentially expressed on various tumor types, blockade of the integrin provides a targeted means of disrupting TGFβ signaling in the TME locally while avoiding the off-target effects inherent to systemic TGFβ blockade [11]. Of note, αvβ6 also has cell intrinsic effects such as increasing angiogenesis through FAK pathway signaling or promoting chemoresistance through an ITGB6-ERK/MAP kinase pathway [15, 16]. Through its mediation of TGFβ signaling and its cell-intrinsic effects, upregulation of αvβ6 by malignant cells acts upon nearly all cancer hallmarks. Therefore, ITGB6 is a highly attractive therapeutic target that shows much promise for clinical development.

## Materials and Methods

### TCGA RNA-seq analysis

A Human Protein Atlas query surveyed the upregulation of ITGB6 across cancer types (**Figure 1A**). Cancer ITGB6 expression was then compared to expression in paired adjacent normal tissue using the TNMplot tool on a concatenated set of GEO, GTex, TCGA, and TARGET RNA-seq databases that were normalized according to the DESeq2 pipeline [17, 18] (**Figure 1B**).

**Figure 1.**
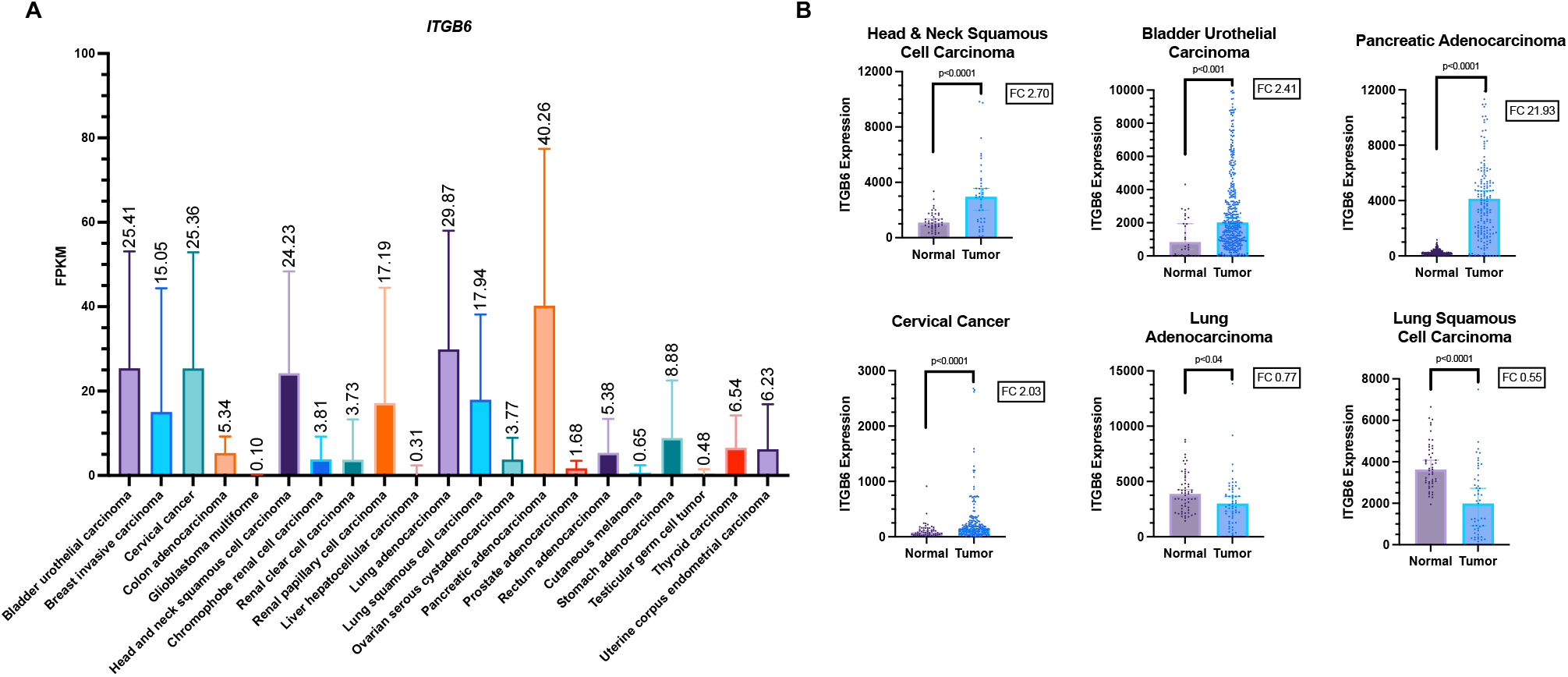
ITGB6 shows specificity and upregulation in human cancers. (A) Human Protein Atlas analysis of RNA expression of ITGB6 across various cancer types. Normalized RNA expression levels are given in fragments per kilobase million (FPKM). Data are acquired from TCGA. Graphs show the mean and SD. (B) Differential gene expression analysis of ITGB6 across tumor and normal tissue. Data are obtained from a standardized concatenated data set from GEO, GTex, TCGA, and TARGET databases. RNA-seq tumor and normal tissue samples are obtained from the same patients and from adjacent sites. Graphs were generated using the TNMplot tool and were normalized using DESeq2. Fold-change (FC) of medians and Mann-Whitney test p-values are shown.

### Cell culture

Both CAL27 and FaDu head and neck squamous cell carcinoma cells were cultured using high glucose DMEM medium with L-glutamine (Cytiva, # SH30022.02) with 10% FBS added. HCT116WT and Capan-2 cells were cultured using high glucose McCoy’s 5A with L-glutamine (Cytiva, # SH30200.FS) and 10% FBS. TALL-104 cells were cultured with RPMI-1640 medium with L-glutamine (Cytiva, cat # SH30027.LS) with 20% FBS and IL-2 (Miltenyi, # 130-097744) added as per ATCC guidelines. The cell lines were kept in a 5% CO2 incubator at 37 degrees Celsius. CT-26, CAL27, FaDu, Capan-2, TALL-104, and HCT116WT were obtained from the American Type Culture Collection. MOC1 and MOC2 cells were obtained from Sigma-Aldrich and KPCY from Kerafast. Human cell lines were authenticated by short tandem repeat profiling. Mycoplasma infection testing was performed on all cell lines.

### Flow Cytometry of Cell Lines

The human cell lines HCT116WT, Capan-2, FaDu, and CAL27 were screened via flow cytometry using an APC-conjugated human ITGB6 antibody (R&D Systems, # FAB4155A). The mouse cell lines CT26, MOC1, MOC2, and KPCY were screened using a mouse/human reactive ITGB6 antibody (Abcam, # ab77906) incubated with an Alexa Fluor 647 conjugated IgG secondary antibody (Thermo Scientific, # A-31571). Cells were gated on singlets and dead cells were excluded using a Zombie Green cell death dye (BioLegend, # 423111).

### CRISPR Knockout of ITGB6

Knockout of ITGB6 was performed by transfection of Alt-R S.p. Cas9-GFP V3 (ID Technology, # 10008100) and single guide RNAs with Lipofectamine CRISPRMAX transfection reagent (Thermo Scientific, # CMAX00003). Single guide RNAs for human ITGB6 were KO-1: GCTAATATTGACACACCTGA and KO-2: CCTGGCTATTCTTCTCATCG. Single Guide RNAs for mouse ITGB6 were KO-1: GCTAATATTGACACACCTGA and KO-2: CGTCATCCATAGAGGCGGAG. The non-targeting sequences was CTRL: GAGCTGGACGGCGACGTAAA.

### Western Blotting

Cells were harvested after CRISPR knockout and lysed in RIPA buffer (Sigma-Aldrich, # R0278) with a protease inhibitor cocktail (Sigma-Aldrich, # 04693159001). Verification of CRISPR knockouts was confirmed using western blot ITGB6 antibodies for human (R&D Systems, # MAB41551) and mouse (R&D Systems, #AF2389) cell lines, respectively.

### Colony Formation Assay

Colony formation assays were performed in 12-well plates over 10 days. CAL27 and FaDu cells were plated at a density of 300 cells per well, while MOC1 and KPCY cells were plated at 200 cells per well. Colonies were counted using ImageJ software if they were greater than 50 cells in size. The average of three replicates was calculated.

### Cancer and immune cell co-culture

The generated CRISPR ITGB6 knockout, and respective control cells of CAL27 and FaDu head and neck squamous cell carcinoma cell lines were fluorescently dyed using CMFDA (Thermo Scientific, # C2925) and plated at a density of 10,000 cells per well in a 96-well plate. After the cells had been allowed to adhere overnight, CMAC (Thermo Scientific, # C2110) fluorescently labeled TALL-104 cells were plated, also at a density of 10,000 cells per well with ten replicates. The experiment contained both co-culture conditions and tumor cells alone for both ITGB6 knockout and control cells. Additionally, the cell death marker Zombie Yellow (BioLegend, # 423103) was added to the culture before imaging. The cells were imaged at 24 hrs at 20X magnification using a Molecular Devices ImageXpress Confocal HT.ai fluorescent plate reader (**Figure 2F, G**). Cell counts of cancer cells as well as the percentage of cancer cells that were dead were quantified using colocalization of the cancer cell and cell death marker fluorescent channels using the Molecular Devices software (**Figure 2H-K**).

**Figure 2.**
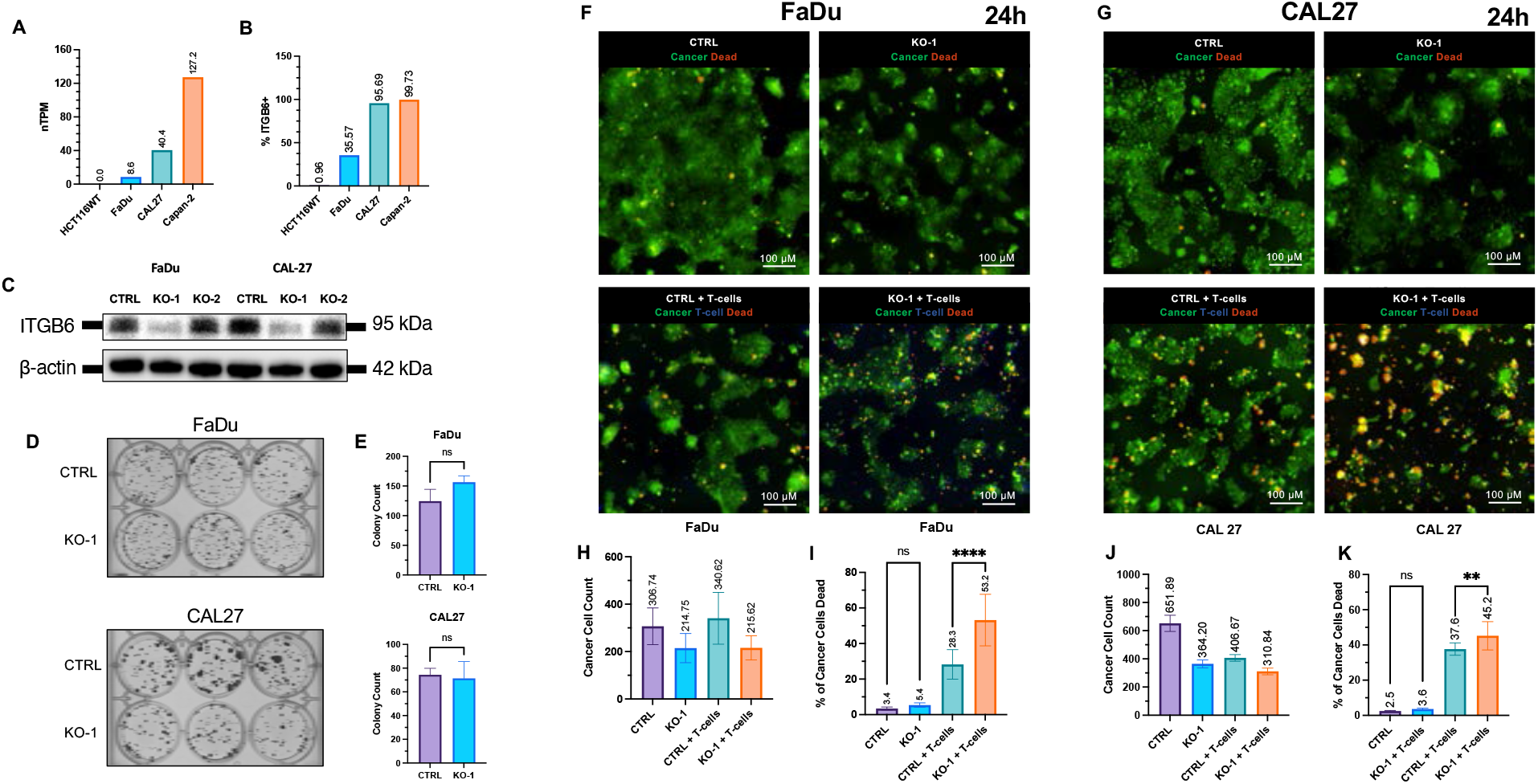
ITGB6 knockout induces T-cell killing of HNSCC cells. (A) ITGB6 RNA of selected human cancer cell lines from the Human Protein Atlas. (B) Flow cytometry analysis of the percentage of cancer cells that are ITGB6+ based on unstained controls. (C) Western blot validation of the CRISPR knockout of ITGB6 in FaDu and CAL27 cells. (D) Colony formation assay of CRISPR control (CTRL) cells and ITGB6 knockouts (KO-1) for FaDu and CAL27 cells. (E) Quantification of colony formation assay. Statistical analysis was performed using an unpaired t-test. Randomly chosen fields of view from the co-culture of FaDu (F) or CAL-27 (G) HNSCC cells (green) and TALL-104 T-cells (blue) (n = 10). Cells were pretreated for 8 hrs with latent TGFβ. Quantification of cancer cell counts for FaDu (H) and CAL27 (J). Quantification of the percentage of cancer cells that are dead for FaDu (I) and CAL27 (K). One-way ANOVA: p < 0.0001 (****), p < 0.002 (**).

### *In Vivo* Studies

C57BL/6 mice were obtained from Jackson Laboratories (Nashville, TN) and were housed in a BSL-2 pathogen-free facility. Mice were 14 weeks old at the beginning of the in vivo study. Mice received subcutaneous flank injections of either 1.5 × 10^6^ MOC1 CTRL or MOC1 ITGB6 KO-1 cells or 1.5 × 10^5^ KPCY CTRL or KPCY ITGB6 KO-1 cells. Cancer cells across the KPCY and MOC1 cohorts were diluted to be volumetrically equal and injected at a 1:1 ratio with Matrigel (Corning, # 356231) for a total injected volume of 200 μL. Each of the four cohorts included ten mice at an equal male-to-female ratio. Tumor volumes were measured starting at day five post-injection (**Figure 3F, H**). Tumor volume was recorded according to the formula,

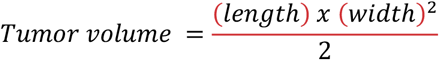

where width is the smaller of the two dimensions. Mice were weighed and their tumor volumes were measured until day 25 post-injection when the mice were euthanized, and their tumors and organs were harvested. Tumors that did not reach a total volume of 100 mm^3^ by the final day were excluded from the analysis. Tumors were surgically removed and gently washed before being weighed (**Figure 3G, I**).

**Figure 3.**
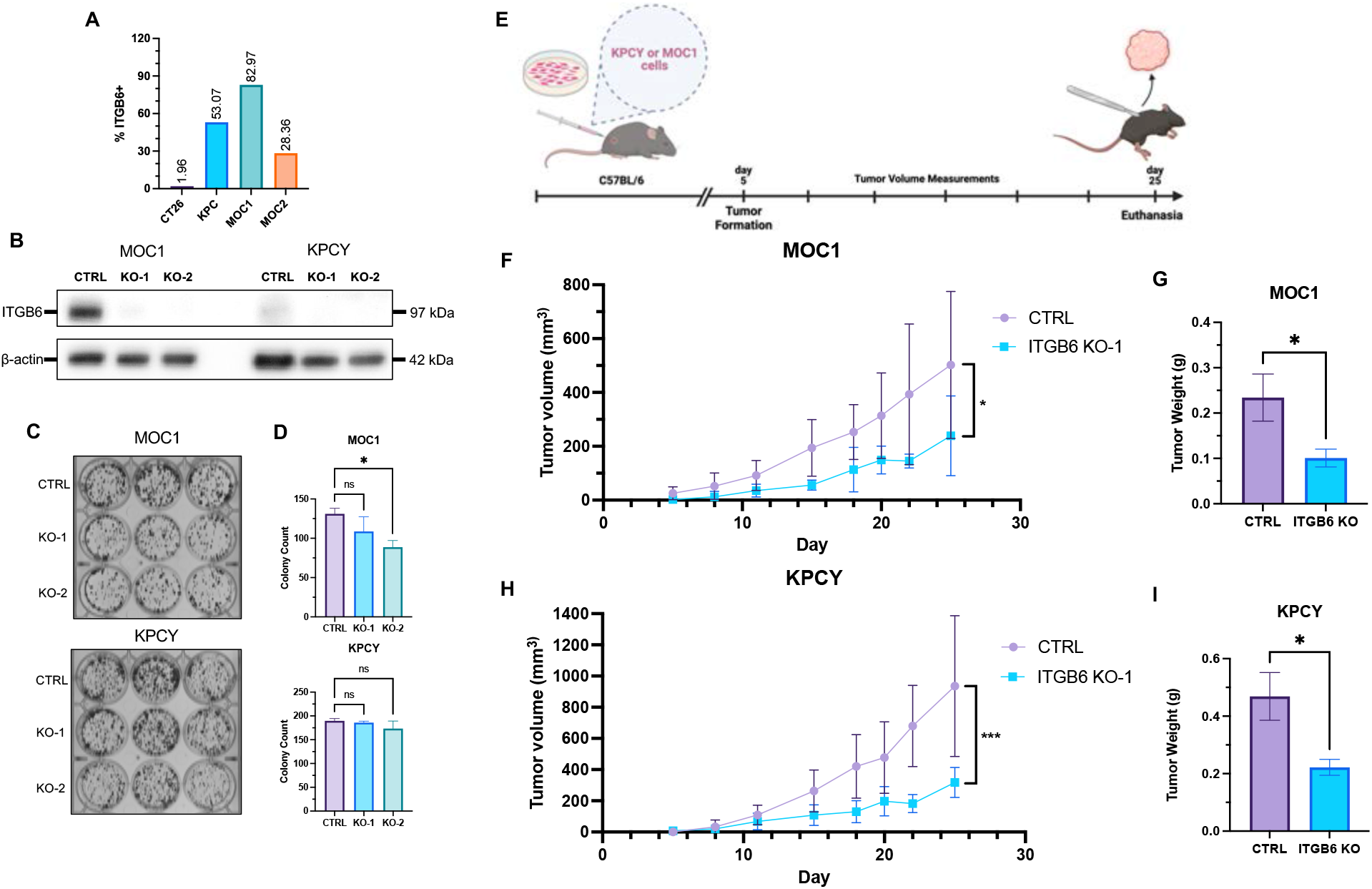
Delayed growth of ITGB6-knockout tumors in immunocompetent mice. (A) Flow cytometry analysis of the percentage of mouse cancer cells that are ITGB6+ based on unstained controls. (B) Western blot validation of CRISPR knockdown of ITGB6 in MOC1 and KPCY cells. (C) Colony formation assay of CRISPR control (CTRL) cells and ITGB6-knockout (KO-1, KO-2) cells for MOC1 and KPCY cells. (D) Quantification of colony formation assay. Statistical analysis was performed using one-way ANOVA: p < 0.05 (*). (E) Experimental timeline for C57BL/6 mice injected with syngeneic tumor cells. Tumor volume of mice starting upon tumor formation at day 5 for MOC1 (F) and KPCY (H) cohorts (n = 10). Mass of MOC1 (G) and KPCY (I) tumors harvested at day 25. Statistical comparison between CTRL and KO-1 cohorts was performed using an unpaired t-test: p < 0.001 (***), p < 0.05 (*).

### RNA-seq analysis of patient treatment and outcome data

Kaplan-Meier curves were generated using TCGA RNA-seq data through the cBio portal, splitting the cohorts according to an ITGB6 mRNA expression threshold of 0.25 standard deviation (σ) above and below the mean for the ITGB6 high and ITGB6 low cohorts, respectively. ITGB6 expression and ICB response were evaluated with the Kaplan-Meier Plotter tool and the ITGB6 expression levels between ICB responders and non-responders were quantified using the ROC Plotter tool [19, 20]. The data was extracted using Python and the figures were generated using Prism.

## Results

### ITGB6 is upregulated in cancer and is specific to malignant tissue

To determine the cancer types potentially susceptible to inhibition of ITGB6, a TCGA analysis of ITGB6 expression was performed (**Figure 1A**). Head and neck squamous cell carcinoma (HNSCC), pancreatic adenocarcinoma (PAAD), cervical cancer, lung adenocarcinoma, squamous cell carcinoma, as well as bladder cancer all demonstrated high RNA expression of *ITGB6*.

The degree of specificity of ITGB6 expression to cancerous tissue versus normal tissue would reveal the efficacy of using blockade of ITGB6 as a targeted therapy against cancer. For the cancer types identified to have high ITGB6 expression, the comparison to the respective native tissues was drawn using RNA-seq data (**Figure 1B**). Head and neck squamous cell carcinoma (median fold-change FC 2.70, p<0.0001), bladder urothelial carcinoma (FC 2.41, p<0.001), pancreatic adenocarcinoma (FC 21.93, p<0.0001), and cervical cancer (FC 2.03, p<0.0001) showed upregulation of ITGB6 compared to normal tissue. Interestingly, lung adenocarcinoma (FC 0.77, p<0.04) and lung squamous cell carcinoma (FC 0.55, p<0.0001) showed downregulation of ITGB6 compared to normal tissue.

### Knockdown of ITGB6 induces T-cell killing of HNSCC cells

After preliminarily screening human cell lines based on their ITGB6 RNA levels, HNSCC cell lines FaDu and CAL27 as well as PAAD cell line Capan-2 were subjected to flow cytometric analysis of ITGB6 expression (**Figure 2A, B**). With the colon carcinoma cell line HCT116WT serving as a negative control (0.96% ITGB6+), FaDu cells showed moderate ITGB6 expression (35.57%), while CAL27 and Capan-2 cells showed high ITGB6 expression (95.69% and 99.73% respectively) (**Figure 2B**).

Furthermore, cell lines FaDu and CAL27 were transfected with Cas9 protein inducing a high-efficiency CRISPR knockout out of ITGB6 with one of the guide RNAs tested. These results were validated via western blot (**Figure 2C**).

The knockout cells with guide RNA-1 (KO-1) as well as the control guide cells (CTRL) were co-cultured with TALL-104 T-cells to study the immunoprotective effect of ITGB6 in HNSCC cells (**Figure 2F, G**). The co-cultures were treated with 10 ng/ml of latent TGFβ for 8 hours to simulate sequestered TGFβ present in the tumor microenvironment. The experiment was conducted with 10 replicates per condition. Quantitative fluorescence microscopy of the co-culture revealed that the knockout of ITGB6 substantially increased T-cell killing over 24 hr in both FaDu cells (FaDu CTRL 28.26% ± 8.29 dead vs. FaDu KO-1 53.18% ± 14.50 dead, p < 0.0001) and CAL27 cells (CAL27 CTRL 37.64% ± 3.51 dead vs. CAL27 KO-1 45.21% ± 8.02 dead, p < 0.01). In the conditions without T-cells, the number of ITGB6 knockout cancer cells was reduced relative to CTRL cells, indicating that the loss of ITGB6 may alter the intrinsic characteristics of cancer cells (**Figure 2H, J**). To account for any intrinsic effects and to measure only T-cell mediated killing of cancer cells, the percentage of dead cancer cells was determined by counting only events of colocalization of the cell death dye (yellow) onto the fluorescent channel of the cancer cells (green) (**Figure 2I, K**). This method allows for the quantification of cancer cell death mediated by cytotoxic T-cells while accounting for any potential ITGB6 status dependent differences in cancer cell growth characteristics. Furthermore, the knockout condition visually exhibited a higher density of T-cells in the proximity of cancer cells as compared to the control condition (**Figure 2F, G**).

Additionally, a 10-day colony formation assay revealed no significant change in cell survival and colony formation potential between control and ITGB6 knockout cells in both models of HNSCC (**Figure 2D, E**). The absence of a differential tumor colony formation rate over 10 days, further suggests that the increased cancer cell death seen over the relatively short 24 hr co-culture was immune-mediated.

### Tumor ITGB6 knockout reduces tumor progression in immunocompetent mice

To demonstrate the anti-tumor activity of ITGB6 knockout *in vivo*, suitable HNSCC and PAAD cell lines were identified for heterotopic injection into immunocompetent mice. Identification of cell lines with high ITGB6 expression was performed via flow cytometry. The CT26 colon carcinoma cell line, known to have low ITGB6 expression, was used as a negative control (1.96% ITGB6+) [21]. The HNSCC cell line MOC2 showed moderate ITGB6 expression (28.36% ITGB6+) while KPCY and MOC1 showed high expression (53.07% and 82.97% respectively) (**Figure 3A**).

As with the human cell lines, MOC1 cells and KPCY were selected for ITGB6 knockout using CRISPR, which was validated via western blot (**Figure 3B**).

To validate that any change of *in vivo* growth rate is not primarily attributed to differences in intrinsic tumor growth characteristics between ITGB6 knockout and control cells, colony formation assays were performed (**Figure 3C, D**). After 10 days, MOC1 cells showed a slightly decreased colony formation rate across both ITGB6 guide RNAs, with knockout cell line KO-2 reaching statistical significance (p < 0.05). KPCY cells showed no observable difference in colony formation rate.

A MOC1 and a KPCY cohort of C57BL/6 mice received flank injections of either CTRL or KO-1 cells (**Figure 3E**). Each of the four groups contained 10 mice. The ITGB6 knockout cells demonstrated reduced tumor progression compared to control cells in both MOC1 (MOC1 CTRL 502.46 mm^3^ ± 273.45 mm^3^ vs. MOC1 KO-1 239.03 mm^3^ ± 147.85 mm^3^, p = 0.0217) and KPCY (KPCY CTRL 935.83 mm^3^ ± 451.08 mm^3^ vs. KPCY KO-1 317.76 mm^3^ ± 95.80 mm^3^, p = 0.0002) cohorts (**Figure 3F, H**). Additionally, there was a substantial difference in the mass of the excised tumors at the end of the experiment for both MOC1 (MOC1 CTRL 0.234 g ± 0.147 g vs. MOC1 KO-1 0.101 g ± 0.059 g, p = 0.0239) and KPCY (KPCY CTRL 0.469 g ± 0.287 g vs. KPCY KO-1 0.222 g ± 0.092 g, p = 0.0128) cohorts (**Figure G, I**).

### High ITGB6 leads to decreased survival in squamous cell cancers and pancreatic adenocarcinoma

TCGA patient outcome data revealed that high ITGB6 expression was a potent marker of a poor prognosis in head and neck squamous cell carcinoma (high ITGB6 35.51 mos. vs. low ITGB6 57.88 mos., p = 0.0090), pancreatic adenocarcinoma (high ITGB6 median survival 15.64 mos., vs. low ITGB6 22.70 mos., p = 0.0002), lung squamous cell carcinoma (high ITGB6 median survival 61.56 mos., vs. low ITGB6 32.85 mos., p = 0.0026), and cervical squamous cell carcinoma (high ITGB6 median survival is undefined, low ITGB6 82.79 mos., p = 0.0222) (**Figure 4**). Statistical analysis was performed using the Log-rank (Mantel-Cox) test.

**Figure 4.**
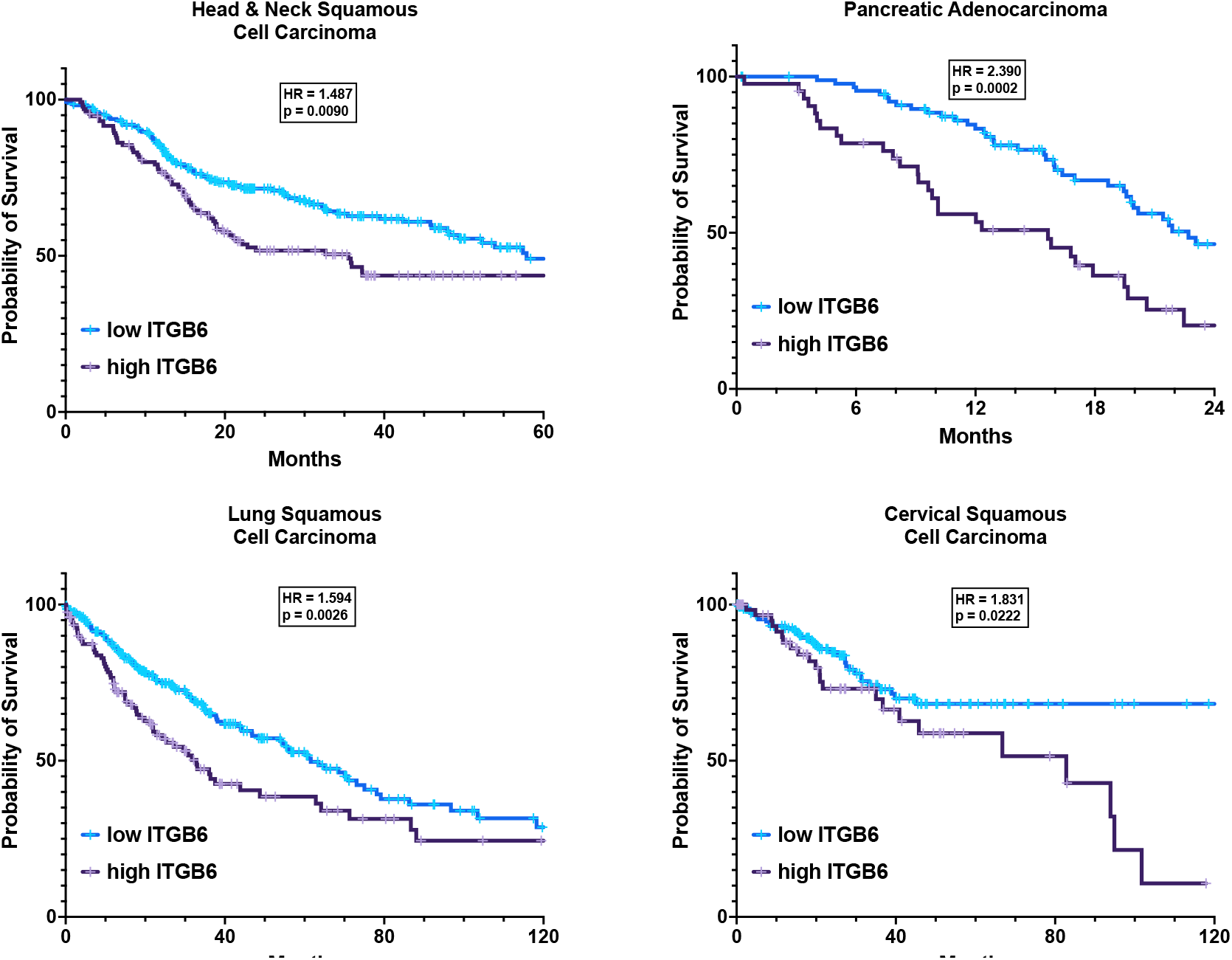
ITGB6 decreases overall survival in patients. Curves show the overall survival of patients with various cancer types stratified based on ITGB6 expression. Patients with high ITGB6 have expression levels above +0.25 σ of the mean and ITGB6 low patients have expression below -0.25 σ. The graphs were generated using TCGA data through the cBioPortal. Statistical analysis of Kaplan-Meier curves and corresponding hazard ratios (HR) was performed using the Log-rank test.

### High ITGB6 expression impairs response to αPD-1 and αPD-L1 immune checkpoint blockade and enhances CTLA-4 response

In a pan-cancer analysis of patients receiving α-PD-1 or α-PD-L1 immune checkpoint blockade, ICB non-responders had higher ITGB6 expression. Non-responders to α-PD-1 therapy had an ITGB6 expression fold change of 1.794 (p = 0.046) while α-PD-L1 non-responders had an ITGB6 fold change of 1.336 (p = 0.023) compared to responders (**Figure 5A**). For patients receiving CTLA-4 therapy, this relationship was reversed, with non-responders having an ITGB6 expression fold change of 0.670 (p = 0.622), however, this relationship did not reach statistical significance.

**Figure 5.**
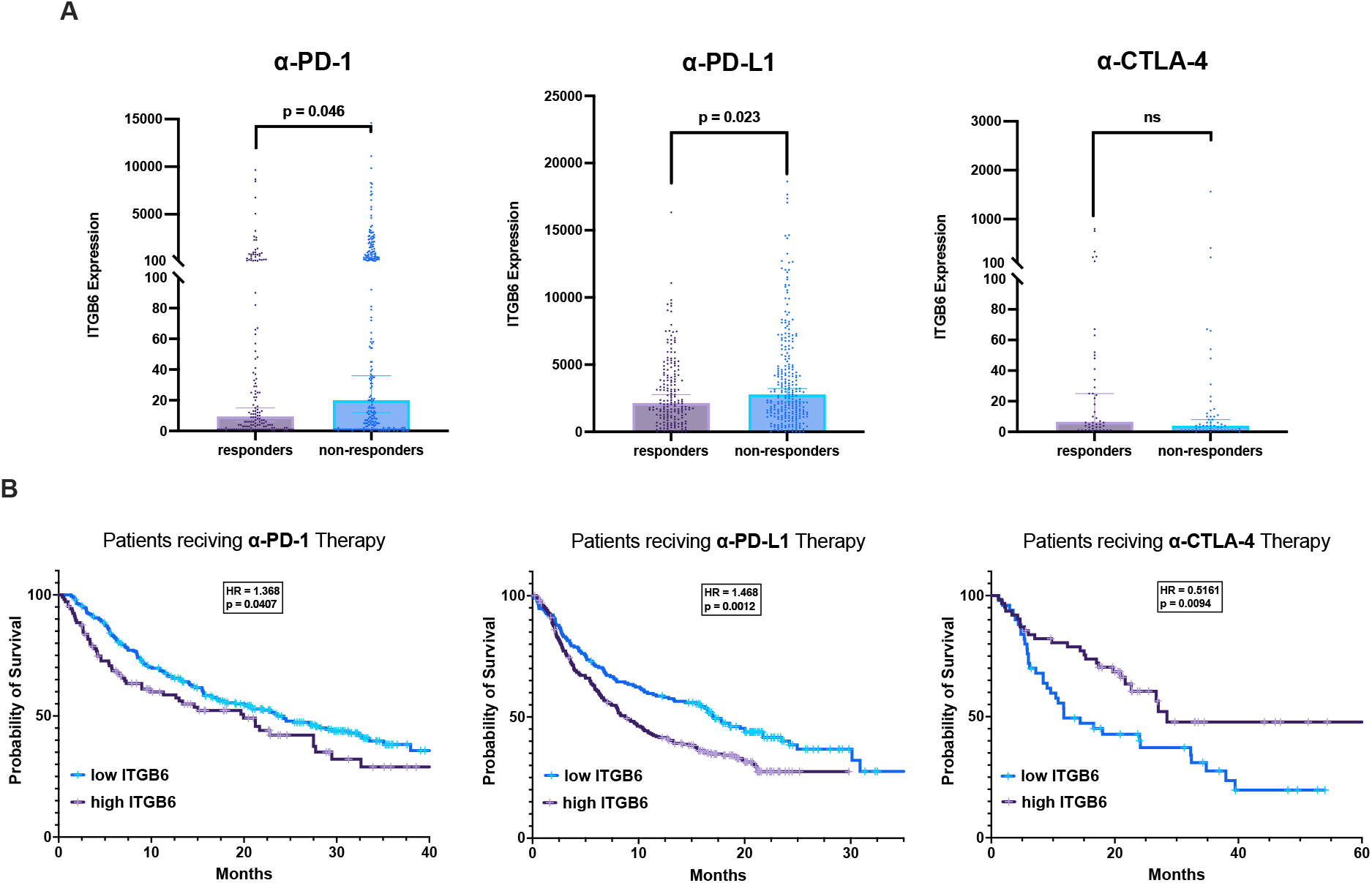
Immune checkpoint blockade is modulated by ITGB6 expression. (A) Pan-cancer analysis of ITGB6 RNA expression level normalized using DESeq2 of patients who are either responsive or unresponsive to immune checkpoint blockade. RNA data are extracted from a TCGA data set of multiple cancer types of patients who underwent immune checkpoint blockade therapy. Graphs were generated using the ROCplot tool and the results of an unpaired t-test are shown. (B) Kaplan-Meier curves of overall survival of patients undergoing immune checkpoint blockade therapy stratified into low and high ITGB6 expression about the mean. Statistical analysis was performed using the Log-rank test.

Furthermore, patient outcome data revealed that high ITGB6 expression was a marker of a poor prognosis in patients receiving α-PD-1 therapy (high ITGB6 19.94 mos. vs. low ITGB6 23.59 mos., p = 0.0407) and patients receiving α-PD-L1 therapy (high ITGB6 8.77 mos. vs. low ITGB6 17.08 mos., p = 0.0012) (**Figure 5B**). However, patients receiving α-CTLA-4 therapy had better outcomes with high ITGB6 expression (high ITGB6 28.43 mos. vs. low ITGB6 11.73 mos., p = 0.0094). Statistical analysis was performed using the Log-rank (Mantel-Cox) test.

## Discussion

The integrin αvβ6, and its β subunit ITGB6, are promising targets for several solid tumors. Squamous cell carcinomas of the head and neck, cervix, and lung, as well as pancreatic adenocarcinomas commonly have upregulated ITGB6 (**Figure 1A**). This upregulation is associated with worse outcomes in patients, including those receiving immunotherapy (**Figure 4, 5B**). Fortunately, the high specificity of ITGB6 to tumor tissue presents as a promising therapeutic opportunity for targeting cancer cells locally while sparing surrounding tissues and minimizing systemic toxicity (**Figure 1B**).

Our in vivo experiments with mouse models of head and neck squamous cell carcinoma (HNSCC) and pancreatic adenocarcinoma (PAAD) showed potent anti-tumor activity in two of the most detrimental cancer types globally (**Figure 3**) [22]. Short timescale experiments with ITGB6 knockout cells, co-cultured with T-cells, indicate that the success observed in our animal models is through immune regulation (**Figure 2**). Additionally, the absence of a substantial growth disadvantage due to ITGB6 knockout in our colony formation assays provides further evidence for an immune-mediated mechanism.

Similar results from αvβ6 blockade in colorectal cancer have convincingly demonstrated a mechanism that attributes the therapeutic effect of αvβ6 blockade to tumor TGFβ paracrine signaling to T-cells using a mouse model with TGFβ receptor-deficient T-cells [21]. Future experiments are planned to further characterize a similar mechanism in HNSCC and PAAD. The patient outcome data correlating ITGB6 expression to immune checkpoint blockade response offers much encouragement to investigate treatment combinations of ITGB6 blockade with clinically relevant immune checkpoint inhibitors such as α-PD-1, α-41-BB, and α-LAG-3. With the efficacy of genetic inhibition of ITGB6 being demonstrated here, future studies can translate the pharmacological and/or biological targeting of ITGB6 into therapeutic clinical trials.

## Acknowledgments

W.S.E-D. is an American Cancer Society Research Professor and is supported by the Mencoff Family University Professorship at Brown University.

